# Perturbation of the insomnia *WDR90* GWAS locus pinpoints rs3752495 as a causal variant influencing distal expression of neighboring gene, *PIG-Q*

**DOI:** 10.1101/2023.08.17.553739

**Authors:** Shilpa Sonti, Sheridan H. Littleton, Matthew C. Pahl, Amber J. Zimmerman, Alessandra Chesi, Justin Palermo, Chiara Lasconi, Elizabeth B. Brown, James A. Pippin, Andrew D. Wells, Fusun Doldur-Balli, Allan I. Pack, Phillip R. Gehrman, Alex C. Keene, S.F.A. Grant

## Abstract

Although genome wide association studies (GWAS) have been crucial for the identification of loci associated with sleep traits and disorders, the method itself does not directly uncover the underlying causal variants and corresponding effector genes. The overwhelming majority of such variants reside in non-coding regions and are therefore presumed to impact the activity of *cis*-regulatory elements, such as enhancers. Our previously reported ‘variant-to-gene mapping’ effort in human induced pluripotent stem cell (iPSC)-derived neural progenitor cells (NPCs), combined with validation in both *Drosophila* and zebrafish, implicated *PIG-Q* as a functionally relevant gene at the insomnia ‘*WDR90*’ locus. However, importantly that effort did not characterize the corresponding underlying causal variant at this GWAS signal. Specifically, our genome-wide ATAC-seq and high-resolution promoter-focused Capture C datasets generated in this cell setting brought our attention to a shortlist of three tightly neighboring single nucleotide polymorphisms (SNPs) in strong linkage disequilibrium in a candidate intronic enhancer region of *WDR90* that contacted the open *PIG-Q* promoter. The objective of this study was to investigate the influence of the proxy SNPs collectively and then individually on *PIG-Q* modulation and to pinpoint the causal “regulatory” variant among the three SNPs. Starting at a gross level perturbation, deletion of the entire region harboring all three SNPs in human iPSC-derived neural progenitor cells via CRISPR-Cas9 editing and subsequent RNA sequencing revealed expression changes in specific *PIG-Q* transcripts. Results from more refined individual luciferase reporter assays for each of the three SNPs in iPSCs revealed that the intronic region with the rs3752495 risk allele induced a ∼2.5-fold increase in luciferase expression (*n*=10). Importantly, rs3752495 also exhibited an allele specific effect, with the risk allele increasing the luciferase expression by ∼2-fold compared to the non-risk allele. In conclusion, our variant-to-function approach and subsequent *in vitro* validation implicates rs3752495 as a causal insomnia risk variant embedded at the *WDR90*-*PIG-Q* locus.

## INTRODUCTION

Sleep is an evolutionarily conserved physiological function experienced across the animal kingdom, including reptiles, amphibians, birds, fish and even jellyfish^1^. Sleep is essential for restoring physical, homeostatic and psychological functions of the body^2^. Conversely, sleep deprivation is known to cause a variety of health problems including, but not limited to, disturbances in cardiovascular, metabolic, and neurological systems. Despite its importance, a substantial proportion of people do not achieve adequate sleep.

Insomnia is a sleep disorder characterized by difficulty initiating or maintaining sleep associated with daytime distress or impairment^3^. Insomnia is common, with 10-15% of the U.S. population meeting criteria for a formal diagnosis and 25-30% experiencing more transient symptoms^4^. Rates of insomnia are higher in women at all ages^5^. Insomnia may occur in relation to factors such as stress, medication side effects, travel, environmental changes, lifestyle and health (physical or psychological) conditions^6,7^. Like other sleep traits, such as sleep duration and chronotype, insomnia is characterized by significant heritability and polygenicity^8,9^.

Genome wide association studies (GWAS) have proven invaluable tools that identify genetic signals contributing to the pathogenesis of a given disease. By comparing the frequencies of alleles in cohorts with and without the trait under investigation, it is possible to detect associated loci for that trait. GWAS successfully revealed risk loci for various sleep traits, typically annotated by the gene residing closest to the association signal. The largest GWAS in the sleep field to date was performed by Janssen *et al*. on insomnia in 1,331,010 participants from the UK Biobank and 23andMe combined, identifying a total of 202 independent risk loci associated with the trait^10,11^.

While GWAS can identify a risk locus, the presence of numerous variants in strong linkage disequilibrium (LD) makes pinpointing the causal variant within that locus challenging. Enrichment of these variants in non-coding (intronic or intergenic) regions further hampers the identification of underlying regulatory variants and the associated risk genes^12^. In addition, the role of *cis*-acting regulatory elements (cREs) that can control genes up to tens or hundreds of kilobases (kb) away further complicates identification of target risk genes^13^.

We developed a “variant-to-gene mapping” strategy that takes proxy SNPs in strong LD with the sentinel GWAS signal that also occupy open chromatin, as determined by ATAC-seq, and then elucidates which of those SNPs reside directly in a region that contacts an open gene promoter using a high-resolution genome-wide promoter-focused capture C approach^14^. Indeed, we have already successfully integrated GWAS summary statistics with our 3D genomic datasets to implicate causal variants and their corresponding causal effector genes for a number of traits, including bone mineral density^14^, systemic lupus erythematosus^15^ and type 2 diabetes^16^.

This approach was applied to the findings from one of the largest insomnia GWAS^10^ to implicate candidate causal genetic variants for this trait. We previously utilized ATAC-seq and promoter-focused Capture C data generated using iPSC-derived neural progenitor cells to identify promoters in open chromatin regions. Insomnia GWAS summary statistics were integrated with 3D genomic data and identified 321 open proxy SNPs corresponding to 100 insomnia loci, of which 36 loci were implicated in an open chromatin loop contacting 88 coding genes^17^ *In vivo* analysis of gene function using RNA interference (RNAi) in *Drosophila* followed by CRISPR-based mutagenesis in zebrafish identified several novel genes with a role in sleep regulation. Of particular note, promoter-focused Capture C in Neural Progenitor cells (NPCs) identified three insomnia-associated proxy SNPs within the *WDR90* locus – rs3752495, rs8062685, and rs9932282 at Chr16p13.3^17^ (r^2^ with sentinel SNP rs3184470 = ∼1) in close proximity with each other, and therefore indistinguishable from each other with respect to possible causality. These SNPs reside across three introns of *WDR90* (rs3752495 – intron 27, rs9932282 – intron 29 and rs8062685 – intron 30) but all looped ∼90 kb to the promoter regions of two candidate effector genes, *NHLRC4* and *PIG-Q*, plus the *NME4* promoter 258 kb away^17^. The presence of a highly conserved ortholog enabled testing the involvement of *PIG-Q* in sleep using two model organisms – fruit fly and zebrafish^17^. *NHLRC4* and *NME4* had either no ortholog or ortholog of moderate/low rank in these organisms, hence were not prioritized for testing. *PIG-Q (*Phosphatidylinositol glycan, Class Q) demonstrated a strong sleep phenotype in both model organisms and was therefore implicated as an effector gene at this insomnia locus^17^. *PIG-Q* is involved in the first step in glycosylphosphatidylinositol (GPI)-anchor biosynthesis that serves to anchor proteins to the cell surface. This gene encodes a N-acetylglucosaminyl transferase that is part of the complex that catalyzes transfer of N-acetylglucosamine (GlcNAc) from UDP-GlcNAc to phosphatidylinositol (PI)^18^.

Given the previous study only focused on the effector gene, *PIG-Q,* there remained a need to characterize the corresponding underlying causal variant at this GWAS signal. The objective of this study was therefore to first identify if CRISPR deletion of the putative regulatory element harboring all three proxy SNPs in human NPCs affects the expression of *PIG-Q* at the gross level, and then to leverage discrete luciferase assays to specifically pinpoint which SNP(s) is a causal variant driving changes in distal gene (*PIG-Q*) expression.

## METHODS

### Cell culture

Human patient derived CHOP10.2P wildtype iPSCs on mouse embryonic fibroblasts (MEFs) were obtained from the Human Pluripotent Stem Cell Core at the Children’s Hospital of Philadelphia. The iPSCs were transitioned into feeder-free conditions inhouse and maintained as feeder-free cultures in mTeSR Plus basal medium supplemented with mTeSR Plus 5X supplement (StemCell Technologies). The cultures were grown in multiwell plates coated with Matrigel® hESC-Qualified Matrix (Corning) and maintained in an antibiotic-free environment at 37°C with 5% CO_2_. Medium was changed every other day and the cells were passaged as colonies with Versene when >70% confluent.

For luciferase assays, the iPSCs were passaged as single cells using Accutase and plated into Matrigel-coated 24-well plates at a density of 50,000 cells/well with 10 µM ROCK Inhibitor. 24 hours post-seeding, the medium containing ROCK Inhibitor was replaced with fresh mTeSR Plus complete medium.

### CRISPR-CAS9 genome editing

A 2kb region encompassing all three SNPs was deleted using CRISPR-Cas9 in patient-derived human iPSCs generated at the Children’s Hospital of Philadelphia (CHOP10.2P). The knock-out clones were generated using a previously published protocol^19^. Briefly, two guide sequences (FWD: 5’ GGCCCGTGGGGCTCTCATCAAGG 3’ and REV: 5’ GGGCCCAAACGCTGAACGGGTGG 3’) were designed to cut the region encompassing the three SNPs and were annealed to generate a 100 bp double-stranded DNA fragment using Phusion High Fidelity DNA polymerase to generate *WDR90*_gRNA. The *WDR90*_gRNA was ligated into a AflII digested gRNA_Cloning Vector (which was a gift from George Church, Addgene plasmid #41824, http://n2t.net/addgene:41824, RRID: Addgene_41824)^20^ using the NEB Gibson Assembly Kit to generate a *WDR90*_gRNA_Cloning vector.

CHOP10.2P iPSCs were transfected with the *WDR90*_gRNA_Cloning vector and the pCas9_GFP vector (which was a gift from Kiran Musunuru, Addgene plasmid #44719, http://n2t.net/addgene:44719, RRID: Addgene_44719)^21^ using a lipid-mediated process (Lipofectamine STEM) and then sorted for GFP-positive cells 48 hours later. Clones were manually isolated 1 to 2 weeks after plating sorted cells and then screened by PCR to confirm gRNA cleavage. PCR screening provided four validated homozygous knock-out clones. Quality control was performed through *de novo* copy number variation (CNV) analysis and karyotyping. While the karyotype of all four clones was normal, one clone presented with a *de novo* CNV (1Mb deletion in 18q23) and was discontinued from further analysis.

### Differentiation to NPCs

Unedited and *WDR90* knock-out iPSC clones were transitioned into feeder-free condition in mTeSR™ Plus medium (STEMCELL Technologies) in a matrigel-coated 6 well plate at a density of 5x10^5^ cells/well with 5 µM ROCK Inhibitor. 24 hours later, the medium was switched to mTeSR Plus medium with two inhibitors (IWR1 – a tankyrase Wnt/β-catenin signaling pathway inhibitor, 1.5μM and SB4315242 – TGF-β receptor kinase inhibitor, 10 μM, Selleck) to induce neuronal differentiation. The medium was replaced every day. On Day 8, the medium was changed to neural expansion medium (1:1:0.04 Neurobasal medium: Advanced DMEM/F12: Neural induction supplement; ThermoFisher). Cultures were maintained in the neural expansion medium until Day 14 and NPCs were harvested. A quality check was performed by measuring NPC-specific markers using flow cytometry. RNA was extracted from Day 14 NPCs (unedited and *WDR90* knock-out) and used for synthesizing libraries for RNA sequencing (RNA-seq) to identify changes in gene expression that occurred at the genome level when the *WDR90* putative enhancer region with proxy SNPs was knocked out.

### RNA-seq library preparation

Day14 NPCs were harvested and pelleted in DNA/RNA shield (Zymo) and stored at -80°C. Total RNA was extracted using the DNA/RNA Miniprep Plus Kit (Zymo), following manufacturer’s protocol. The quality of the RNA was assessed using Agilent 2100 Bioanalyzer and the RNA 6000 nano Assay chip. Samples with high-quality RNA (RNA integrity number (RIN) >7) were used for library preparation. The cDNA libraries were constructed from 1 ug input RNA using the NEBNext® Ultra™ II Directional RNA Library Prep Kit for Illumina® (NEB), following manufacturer’s recommendations. The final cDNA libraries were assessed using Agilent 2100 Bioanalyzer and the DNA 1000 Assay chip. Libraries were sequenced on the Illumina NovaSeq 6000 platform (51 bp read 1 and read 2 length).

### RNA-seq gene-level analysis

Sequencing data was demultiplexed to generate FASTQ files using Illumina bcl2fastq2 Conversion Software. FASTQ files were assessed with FastQC^22,23^ to verify that there was high sequence quality, expected sequence length, and no adapter contamination. Paired-end FASTQ files for each replicate were mapped to the Ensembl human reference transcriptome (GRCh38)^24^ using Kallisto^25,26^. Abundance data generated with Kallisto was read into R, annotated with Ensembl human gene annotation data (version 86)^24^, and summarized as counts per million (cpm) at the gene level. Genes with less than 1 cpm in 3 samples were removed to increase statistical power to detect differentially expressed genes. Samples were normalized with the trimmed mean of M values (TMM) method^27^. The R package ‘limma’^28^ was used to identify differentially expressed genes by first applying precision weights to each gene based on its mean-variance relationship using the voom function and then linear modeling and Bayesian statistics were employed to detect genes that were up- or down-regulated in each condition. Genes with an adjusted P-value < 0.05 and |log2 fold change| > 1 were considered significantly differentially expressed. Significantly differentially expressed genes were clustered using Pearson correlation and the R function hclust. The clustered genes were cut into 2 modules and significantly enriched Gene Ontology terms^29,30^ in each module were identified using the R package ‘gprofiler2’^31,32^.

### RNA-seq transcript-level analysis

Sequencing data was demultiplexed to generate FASTQ files using Illumina bcl2fastq2 Conversion Software. FASTQ files were assessed with FastQC^22,23^ to verify that there was high sequence quality, expected sequence length, and no adapter contamination. Paired-end FASTQ files for each replicate were mapped to the Ensembl human reference transcriptome (GRCh38)^24^ using Kallisto^25,26^. Abundance data generated with Kallisto was read into R, annotated with Ensembl human gene annotation data (version 86)^24^, and summarized as log2 counts per million (cpm) at the transcript level. Transcripts with greater than 0.2488206 cpm in at least 3 samples were retained after filtering. This threshold was chosen because it selected for transcripts with at least 10 counts in the smallest library sample. Samples were normalized with the trimmed mean of M values (TMM) method^27^. The R package ‘limma’^28^ was used to identify differentially expressed transcripts by first applying precision weights to each transcript based on its mean-variance relationship using the VOOM function and then linear modeling and Bayesian statistics were employed to detect transcripts that were up- or down-regulated in each condition. Transcripts with an adjusted *P*-value < 0.05 and |log2 fold change| > 1 were considered significantly differentially expressed.

### Quantitative real-time PCR validation

The selected primers used for qRT-PCR were designed using Primer 3.0 and synthesized by IDT. β-actin was used as an internal control for all samples. Primer sequences were as follows: *WDR90*-CRISPR forward, 5’-ACT CTG CCA AGG GCA CTT GCC -3’ and reverse, 5’-GGC CAG GAC CTG GGC ACT G -3’; *WDR90* forward, 5’-CTG GAG GCA GAG CAC GAG GG -3’ and reverse, 5’-AGT CGT ATA GCT GCT GCA GGG T -3’. qRT-PCR was performed using the AriaMx Real-time PCR System with a PowerUP SYBR green expression assay system. The PCR reaction conditions were as follows: UDG activation at 50°C for 2 minutes, followed by Dual-lock DNA polymerase activation at 95°C for 2 minutes, followed by 40 PCR cycles of denaturing at 95°C for 15 sec and annealing and extension 60°C for 60 sec (manufacturer’s protocol for primers with Tm > 60°C). Each sample was assayed in triplicate. The 2-ΔΔCt method was used to determine fold-change in gene expression in the *WDR90* KO NPC samples relative to the unmodified WT NPC samples. For statistical analysis, we used one way ANOVA to compare the expression of *WDR90* between *WDR90* KO and WT samples. *P*<0.05 was considered to indicate a statistically significant difference.

### Functional group analysis

The Gene Ontology (GO) program provides “terms” that can describe the function of a gene in any organism (http://www.geneontology.org)^29,30^. These terms can be broadly categorized into: Biological process (BP), cellular component (CC) and molecular function (MF). The *P*-value was used to denote the significance of the GO term enrichment in the module of differentially expressed genes, where enrichment was considered significant if *P*<0.05.

### Splicing Event analysis

The software package SplAdder(v3.0.4)^33^ was used to detect splicing alterations by identifying events found in annotated transcripts and detect novel splice events. Spladder builds and augments splice graphs and subsequently quantifies splice events. GencodeV30 was used as the initial gene reference (https://ftp.ebi.ac.uk/pub/databases/gencode/Gencode_human/release_30/). Reads were aligned to hg38 using STAR (v2.7.9) and indexed using samtools index (v1.11). The various events considered by this package to detect splicing were exon skipping, intron retention, alternative 3’ splice sites (A3SS), alternative 5’ splice sites (A5SS) and mutually exclusive exon (MXE). Splice graphs were built for individual replicates, and subsequentially merged using the spladder build with parameters --no-extract-ase --ignore-mismatches, and either --merge-strat merge_graphs for merged graphs or --merge-strat single for individual replicates. Plots were visualized with ggplot2 (v)^34^. Events were quantified using spladder build with the parameters --merge-strat merge_graphs --no-extract-ase --quantify-graph --qmode single --ignore-mismatches. Spladder test was performed with the default settings, which models junction read count with a general linear model framework for differential testing, as performed by standard differential testing approaches like DEseq2. Read count was adjusted per replicate from estimating the mean-variance relationship to account for overdispersion.

### Cloning

A putative enhancer region encompassing approximately 550 bp on either side of each proxy SNP was identified and amplified using PCR for infusion cloning (Takara), a ligase-free method to clone any insert into any vector. A pGL3-promoter vector was used for performing luciferase assays. pGL3 was linearized using Kpn1 and Not1 restriction enzymes. The primers to amplify the enhancer fragment were designed to be 33-36 bp with at least 15 bp extensions complimentary to the vector ends with restriction overhangs of Kpn1 and Not1, and another 15 bases specific to the enhancer region being amplified and were ordered through IDT. Oligos used for amplifying the enhancer containing each SNP are as follows:

rs3752495:

forward, 5’-TTTCTCTATCGATAGGCAACTTCATCCCACACCCC-3’

reverse, 5’-TCGAGCCCGGGCTAGGCGCTACCGGGAGAACAGAG-3’

rs8062685:

forward, 5’-TTTCTCTATCGATAGTTGTCAGATCCGCGTCTGGG-3’

reverse, 5’-TCGAGCCCGGGCTAGCCAGAGCTGGTGCCACAATAG-3’

rs9932282:

forward, 5’-TTTCTCTATCGATAGGTTCGGCCCCTCCCCAGGCCCCTCCCCGCCCCCCCCCCCCCCCGG-3’

reverse, 5’-TCGAGCCCGGGCTAGGGCCTCCACCCTCCCTCC-3’.

The amplification of the putative enhancer containing rs9932282 proved challenging given the high GC rich content. We changed the primer design to incorporate as much of the GC rich region as we could into one oligo, resulting in a 60 bp oligo.

Following infusion cloning, the recombinant DNA was transformed into NEB Stable *E. coli* cells (NEB) under standard transformation conditions. Colonies were selected and plasmid DNA was extracted using The PureLink™ Quick Plasmid Miniprep Kit (Thermofisher Scientific) and sent for Sanger sequencing to identify the integrity and quality of the inserted enhancer fragment. Vectors with the correct sequence were purified using the PureLink Expi Endotoxin-Free Maxi Plasmid Purification Kit (Thermofisher Scientific) and the sequence was confirmed again using Sanger sequencing. Once the putative enhancers containing all three SNPs were successfully cloned into the pGL3 vector, site directed mutagenesis using the QuikChange II Site-Directed Mutagenesis Kit (Agilent) and the NEB Q5 Site Directed Mutagenesis kit (NEB) was performed to introduce the risk allele for each SNP. The primers used for mutagenesis are as follows:

rs3752495:

forward, 5’-CTCCCACACCTCCCACCCCAGCTTCCC-3’

reverse, 5’-GGGAAGCTGGGGTGGGAGGTGTGGGAG-3

rs8062685:

forward,5’-CTGACGTGGCTGCTGTGTGTCCTTCCC-3’

reverse, 5’-GGGAAGGACACACAGCAGCCACGTCAG-3’

rs9932282:

forward, 5’-CTCACGCCTGCCCTCTTGCCTGC-3’

reverse, 5’-TGGGGAGGCCGTGGCTGG-3’

Following transformation, clones were selected, and the plasmid DNA was Sanger sequenced to confirm clones with the correct sequences. Vectors with the correct sequence were purified using the PureLink Expi Endotoxin-Free Maxi Plasmid Purification Kit (Thermofisher Scientific) and the sequence was confirmed again using Sanger sequencing. The vector DNA with either the risk or non-risk allele was stored at -20°C until the iPSCs were ready to be transfected.

### Transfection

The iPSCs were passed with Accutase into a single cell suspension and seeded in Matrigel-coated 24-well plates at a density of 50,000 cells/well in 2 mL mTeSR Plus medium with 10 µM ROCK Inhibitor to prevent cell death. 24 hours following the seeding, the medium was replaced with fresh mTeSR Plus medium, and the cells were transfected with 500 ng of plasmid DNA (unmodified pGL3-promoter vector, modified pGL3 with each putative enhancer containing each SNP and Renilla luciferase vector (pRL-TK) to serve as an internal control at a ratio of 20:1, respectively) using Lipofectamine LTX with PLUS Reagent. The cells were harvested for the luciferase assay 48 hours later.

### Luciferase assay

48 hours following the transfection of iPSCs, growth media was removed, and cells were washed gently with phosphate buffered saline (PBS). 5X Passive Lysis Buffer (Promega) was diluted to 1X with PBS and 100 µL was added per well to harvest cell lysates. The plate was incubated at room temperature for 30 minutes and with gentle rocking before the cell lysates were harvested for luciferase assay. The fluorescence of firefly and renilla luciferase were measured using the Dual-Luciferase Reporter Assay System (Dual-Glo Luciferase Assay System, Promega) according to the manufacturer’s instructions and using the SpectraMax-iD5 plate reader (Molecular Devices). 20 µL of cell lysate per sample was transferred into a white opaque 96-well plate in triplicate. 100 µL of the Luciferase Assay Reagent II was set to be dispensed into each well by an injector after which there would be a 2-second pre-measurement delay, followed by a 10-second measurement period to measure the fluorescence from firefly luciferase. Then, 100 µL of the Stop & Glo Reagent was dispensed into the same well by a second injector. After a 2-second pre-measurement delay, fluorescence from renilla luciferase was measured for a 10-second measurement period. The fluorescence units obtained for each putative enhancer vector with risk or non-risk alleles and the promoter only vector was first normalized to plate blank, then normalized with the internal control renilla luciferase signal and finally with the promoter only plasmid construct normalized signal. Ten independent experiments were performed. Quantitative data of the reporter gene assay are calculated as mean ± SEM. One-way ANOVA with Tukey’s HSD posthoc test was used to determine significant differences between putative enhancer vectors.

## RESULTS

### Generation and characterization of WDR90 KO NPCs

CRISPR deletion of the putative enhancer region harboring all three SNPs in iPSCs generated three knock-out clones – KO CL 7.2, KO CL 45.1 and KO CL 48.2 – were used for further characterization. The deletion in the *WDR90* putative enhancer region in all three knock-out iPSC clones was confirmed using PCR and qPCR. Unedited and knock-out iPSCs were then differentiated to NPCs (See **Methods**). We also measured *WDR90* expression with qPCR in the knock-out NPCs, and CRISPR deletion decreased, but did not eliminate, the expression of *WDR90* (**Suppl. Fig. 1**).

Following this confirmation, we used RNA sequencing to characterize any differences in global gene expression between the unedited and knock-out NPCs. Perturbation of the genomic region harboring all three SNPs resulted in three main observations: 1: Differences were observed in *PIG-Q* expression at the transcript level; 2: One other PIG family gene was differentially expressed in knock-out cells; 3: Genes involved in the WNT signaling pathway were found to be most differentially expressed – as outlined further below.

### Differences in PIG-Q transcript expressions levels

To characterize the global changes in expression resulting from deletion of these SNPs, we performed RNA-seq on unedited NPCs and the three knockout clones (**Supp. Table 1**). Principal component analysis (PCA) was performed to explore RNA-seq library reproducibility. The first principal component mapped to differences between the wild type and the three knock- out NPC clones (accounting for 49.8% of variance). Technical replicates of each condition largely clustered together (**Fig. 1**). To identify differentially expressed genes, precision weights were first applied to each gene based on its mean-variance relationship using VOOM^35^, then data was normalized using the TMM method^27^ in edgeR^36^. Linear modeling and Bayesian statistics were employed via Limma^28^ to identify genes that were up- or down-regulated in each experimental condition. A total of 1,771 genes were significantly differentially expressed using the criteria of adjusted *P*-value < 0.05 and absolute logFC > 1 (**Fig. 2**).

**Figure 1:**
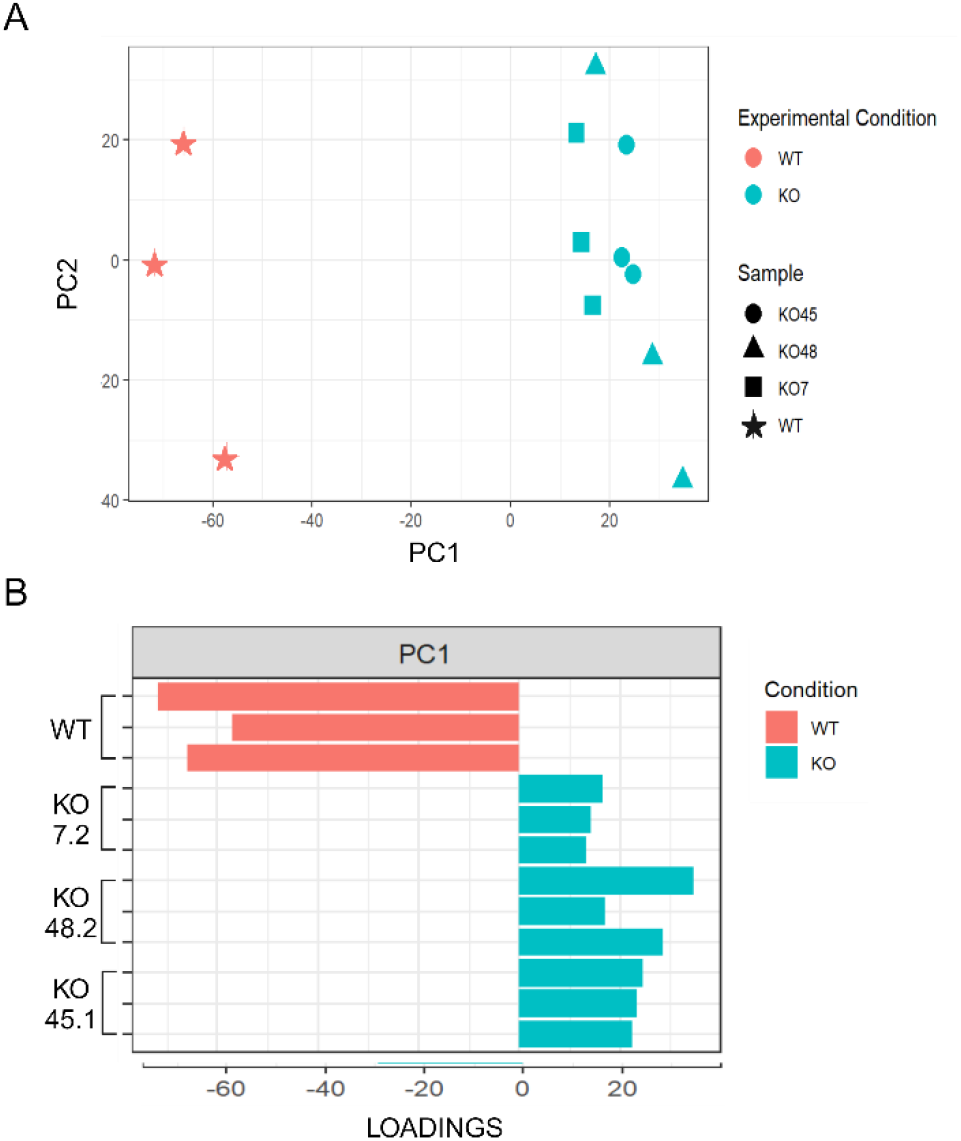
CRISPR edited WDR90 KOs clustered differently from unedited wild type NPCs. A. Principal components analysis (PCA) plot projecting the differences between samples. B. Small multiples plot showing the contribution of each sample to its first principal component (PC1). The x-axis represents loadings. Loadings quantify the contribution of a sample to a particular component. The sign of a loading indicates either a positive or negative correlation. Technical replicates showed similar clustering and contribution to principal components.

**Figure 2:**
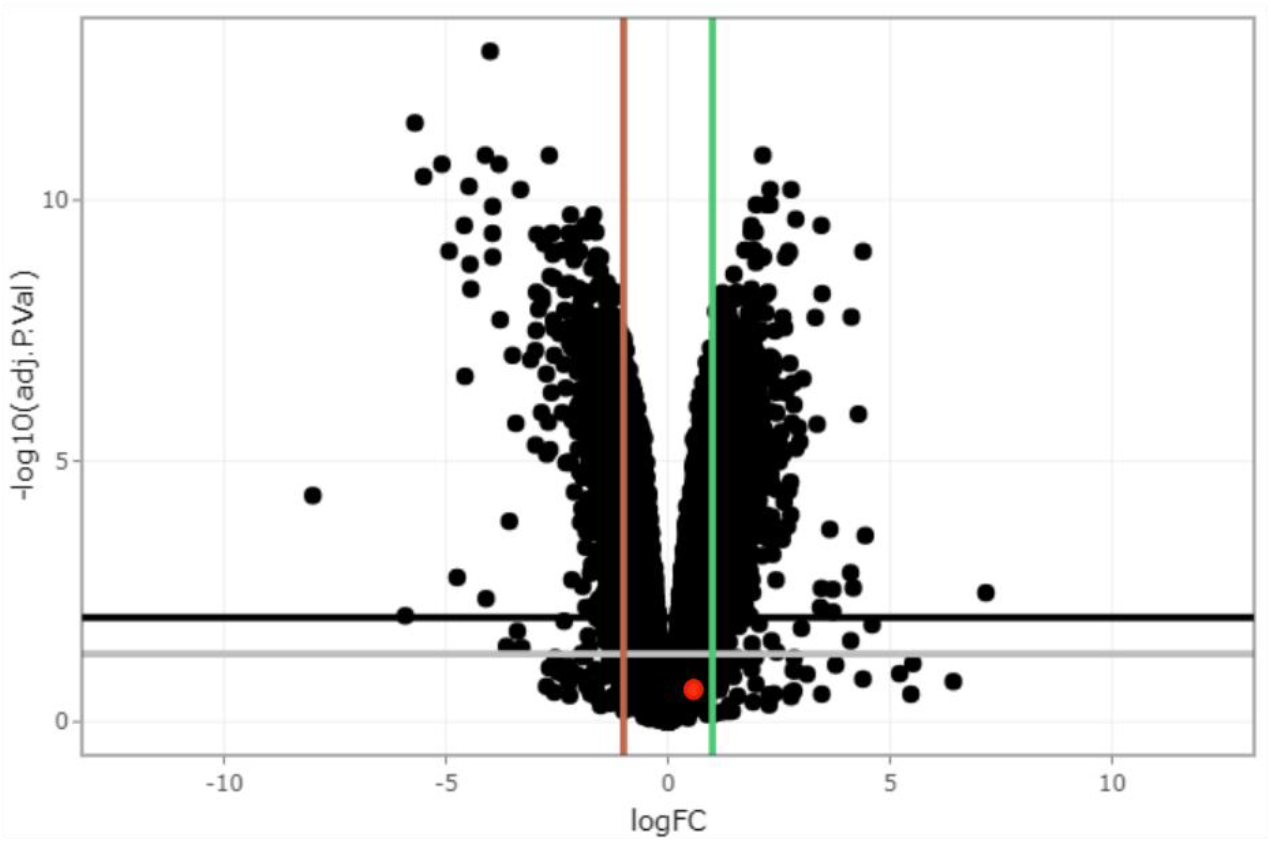
Volcano plot visualizing differentially expressed genes. Data points above the grey line represent genes from the differential expression analysis that had an adjusted *P*-value < 0.05 and data points above the black line were genes that had an adjusted *P*-value < 0.01. Data points to the left of the red line were genes from the differential expression analysis that had a logFC < -1, representing down-regulated genes, and data points to the right of the green line were genes with a logFC > 1, representing up-regulated genes. Red dot represents *PIG-Q*.

Notably, we did not detect *PIG-Q* among the significantly up- or down-regulated genes in the knock-out NPCs (logFC = 0.21; adjusted *P*-value = 0.28) (**Fig. 2, Suppl. Table 2**). However, as many as 25 *PIG-Q* transcript variants have been annotated, of which 9 do not code for protein, 8 are protein coding and 8 are likely to be degraded by nonsense mediated decay (NMD)^24^ (**Suppl. Fig. 2**). Of these 25 annotated transcripts, 16 were expressed in the NPCs and the expression levels of *PIG*-Q-208 and *PIG*-Q-218 were found to differ significantly in knock-out NPCs (*PIG*-Q-208: log fold change = 1.07; and PIG-Q-218: log fold change = -7.16) (**Suppl. Table 3**). *PIG*-Q-208 encodes 574 amino acids (∼63 kDa) and *PIG*-Q-218 encodes 65 amino acids. Despite the fact that these transcript variants code for proteins, they are predicted to be involved in nonsense mediated decay (NMD).

### Differential expression of PIG-family genes

*PIG-Q* encodes Phosphatidylinositol Glycan, Class Q, which is a member of the GPI anchor biosynthesis pathway. This pathway has been previously explored in detail and is reported to involve several other phosphatidylinositol glycan (PIG) genes^37^. In addition to *PIG-Q*, we have previously reported a sleep phenotype in our model organisms for perturbation of other members of the PIG gene family, namely *PIG-Z, PIG-G, PIG-C, PIG-M, PIG-L, PIG-O, PIG-Wb, GAA1* and *PGAP2* ^17^. In this study, knock-out NPCs also significantly differentially expressed *PIG-W* (log fold change = -1.09, adjusted *P*-value = 7.51x10^-6^) (**Suppl. Table 4)**.

### Modulation of the WNT signaling pathway

The differentially expressed genes in knock-out NPCs (**Suppl. Table 2**) included several genes in the WNT signaling pathway (**Suppl. Table 5**). Three RSPO family genes (*RSPO1*: adjusted *P*-value=1.40x10^-13^; log fold change = -4.00; *RSPO2*: adjusted *P*-value=7.62x10^-8^; log fold change = -2.98; *RSPO3*: adjusted *P*-value=3.31x10^-12^; log fold change = -5.69) were down-regulated and encode proteins that function as ligands for leucine-rich repeat-containing G-protein coupled receptors (LGR proteins) and positively regulate the WNT signaling pathway. Other components involved in the WNT signaling pathway that were down-regulated in knock-out NPCs included *WNT4*, *WNT5B*, *WNT7A*, *WNT7B*, *WNT8B* (encoding WNT proteins), *CTNNB1* (encoding beta-catenin), *FZD1* (Frizzled protein), *LEF1*, and *TCF19* (transcription factors). Meanwhile, *FZD8* and *FZD10* were up-regulated in the knock-out NPCs.

### Functional enrichment analysis

Gene Ontology (GO) enrichment analysis identified that biological processes of cell communication, cell signaling and were upregulated in knock-out NPCs. Biological processes such as neurogenesis and neuronal differentiation were down-regulated in the knock-out NPCs, suggesting a key role for *WDR90* and/or *PIG-Q* in modulating pathways implicated in neuronal differentiation (**Fig. 3**).

**Figure 3:**
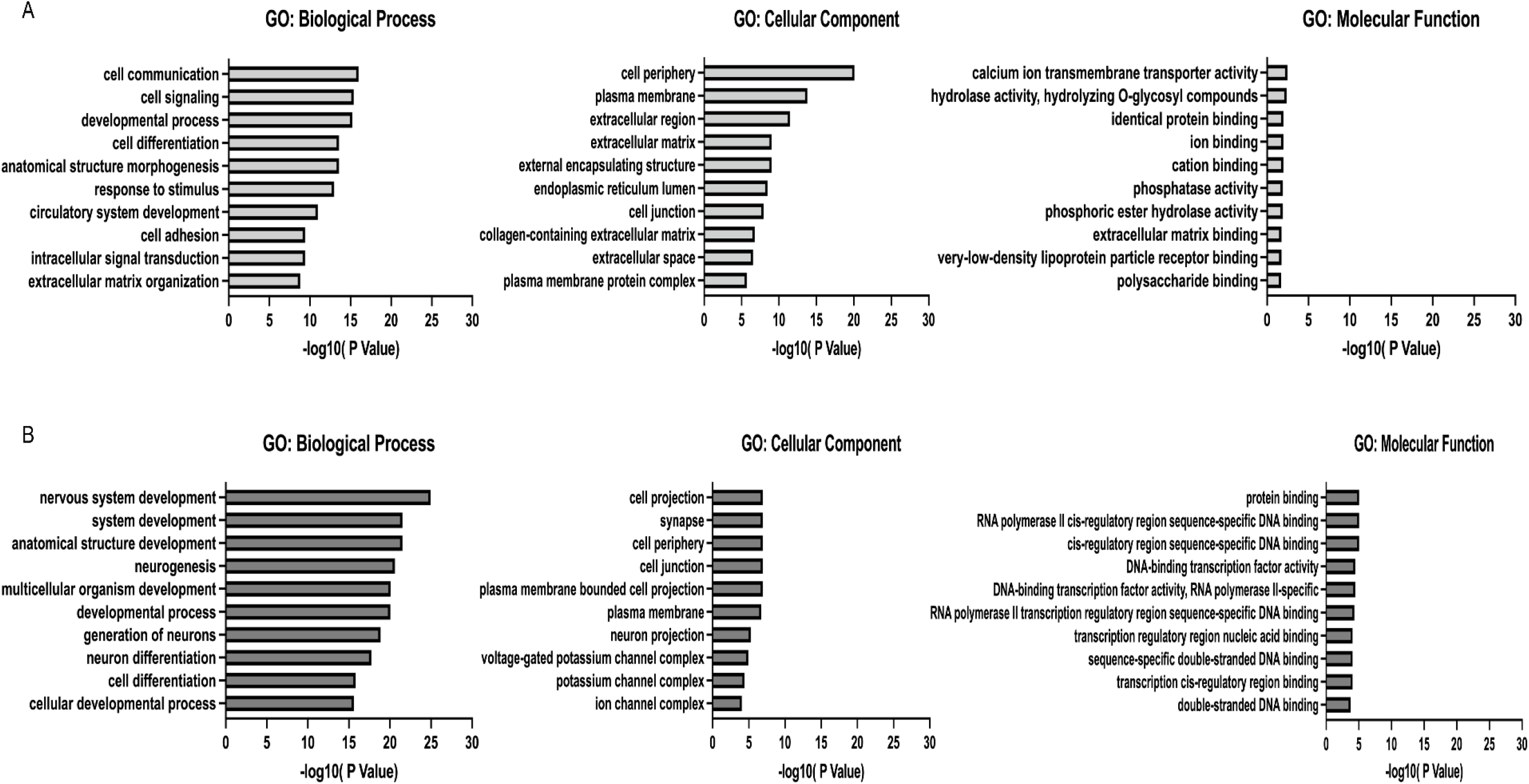
Gene ontology (GO) enrichment analysis. The top 10 enriched GO terms in genes up-regulated (A) and down-regulated genes (B) in KO NPCs. Enrichment score indicates the enrichment score value of the GO term, equivalent to -log10(P-value).

### WDR90 KO modulates splicing in PIG-Q

*PIG-Q* is reported to have 25 splice variants^24^ (**Suppl. Fig. 2**). RNA sequencing can identify the most expressed transcript variant of a gene by estimating which transcript is likely expressed in each experimental condition. We elected to further explore the transcript level differences in *PIG-Q* that we observed in our transcript level analyses with Kallisto^25,26^ and edgeR^36^. The prior analysis depends on the reference transcript annotations, which are limited compared to the transcriptional sequence variation that has been detected across large-scale NGS datasets^38^, and as such alternative approaches allow for quantification of alternate splicing by counting “splicing events” to uncover cryptic exons/splicing events. To this end, we reanalyzed the unedited and knock-out RNA-seq data using the event-based quantification tool, SplAdder^33^. Consistent with expectations of known splicing differences across cells, we identified more events representing *de novo* intron inclusion across comparisons compared to other events (**Fig. 4, Suppl. Table 5**). Examining *PIG-Q*, a cryptic exon was predicted to located within an intron in our NPC data (**Fig. 4B**). This exon displayed a significantly higher probability of splicing inclusion (PSI) in two of the three mutants (KO CL 45.1: dPSI = 0.068, log fold change = 1.82, FDR = 3.32 x10^-4^ and KO CL 48.2: dPSI = 0.063, log fold change = 1.70, FDR = 7.54 x10^-3^). KO CL 7.2 also shown an increased inclusion but did not reach statistical significance (**Fig. 4C**). The exon was well supported with average of ∼20 junction reads in the control NPCs and ∼70 in each mutant. In combination with the differential expression of *PIG-Q* transcripts, this result indicates that deletion of three variants within *WDR90* influences alternative splicing of *PIG-Q*.

**Figure 4:**
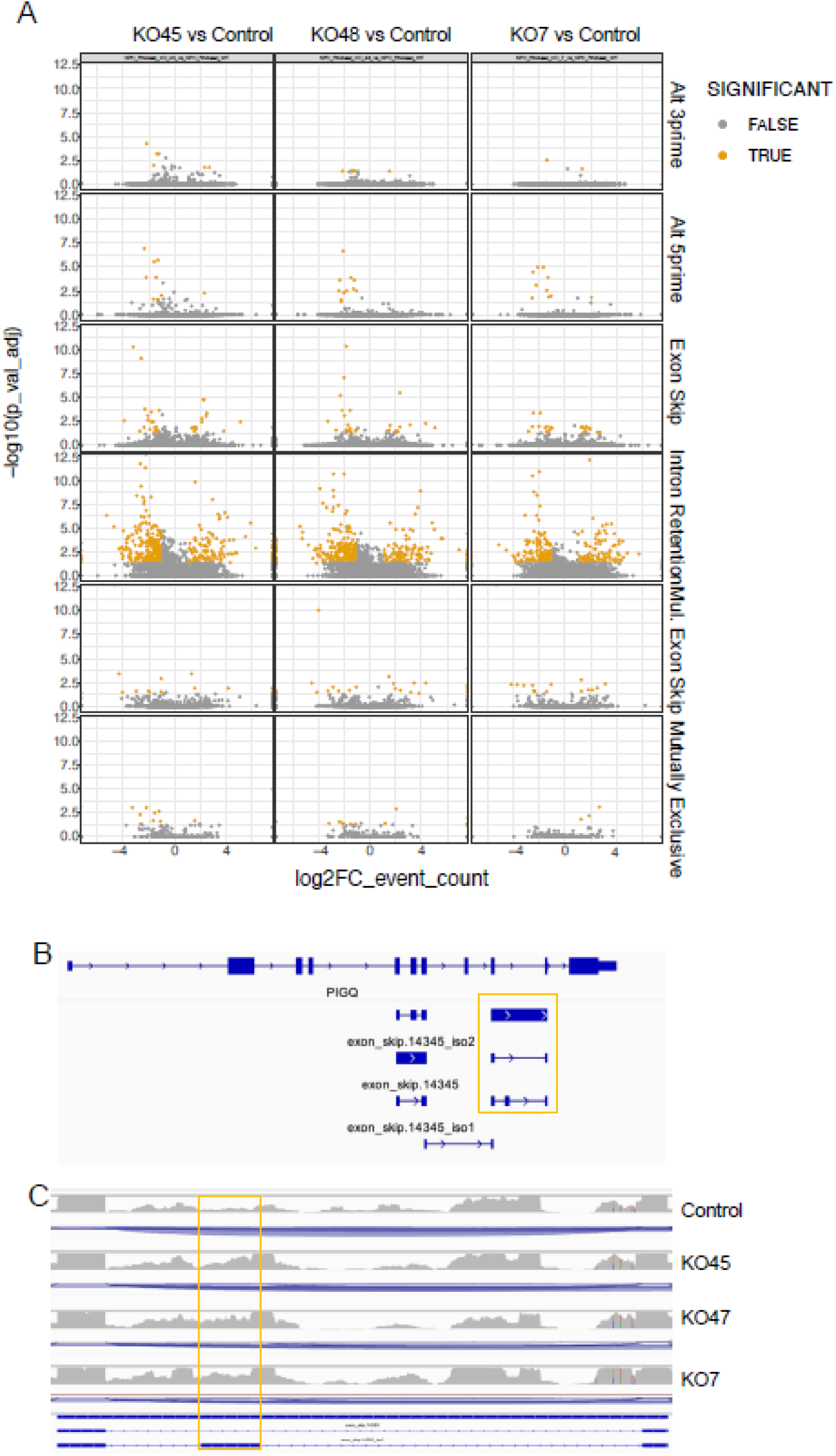
Global splicing comparison of each mutant to control NPCs. Volcano plot showing differential splicing events for each event class. X-axis: log (FC) event count, Y-axis: log (adj p. value).

### Identification of a causal regulatory variant

Given our observation that perturbation of the putative regulatory region harboring the three proxy SNPs influences *PIG-Q* transcript expression, we were therefore motivated to subsequently pinpoint the causal variant(s) among these three variants with luciferase reporter assays. We cloned the ∼550 bp putative enhancer region surrounding each of the three proxy SNPs with either the risk allele (the allele in LD with sentinel SNP, rs3184470, allele associated with risk of insomnia) or the non-risk allele into the pGL3 firefly luciferase vector with a SV40 promoter. We transfected these pGL3 promoter vectors into iPSCs and co-transfected with pRL-TK renilla luciferase vector for a normalization control. Results from luciferase reporter assays for all three SNPs individually revealed that the intronic region with the rs3752495 risk allele exhibited strong enhancer activity, which induced a ∼2.5-fold increase in luciferase expression (*n*=10; *P*<0.05) relative to unmodified pGL3 (**Fig. 5**). The other two SNPs did not significantly affect luciferase expression in the presence of either risk or non-risk allele. This result supports rs3752495 as conferring regulatory potential in iPSCs and as a causal variant.

**Figure 5:**
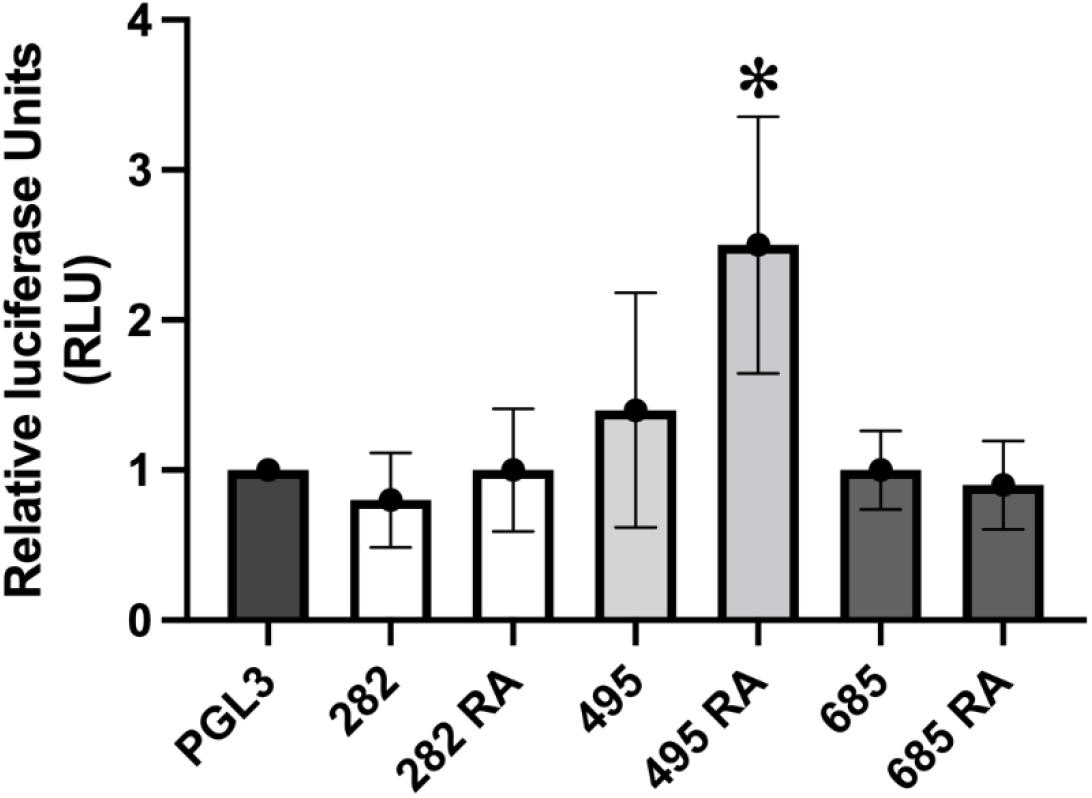
Luciferase reporter assay determines increased regulatory activity with the risk allele of SNP rs3752495. Y-axis represents relative luciferase fluorescence units normalized to renilla internal control and pGL3 promoter only vector. X-axis represents the various experimental conditions – pGL3: promoter only vector, 282: promoter vector with enhancer construct containing non-risk allele of SNP rs9932282 (G), 282 RA: promoter vector with enhancer construct containing risk allele (RA) of SNP rs9932282 (C), 495: promoter vector with enhancer construct containing non-risk allele of SNP rs3752495 (T), 495 RA: promoter vector with enhancer construct containing risk allele (RA) of SNP rs3752495 (C), 685: promoter vector with enhancer construct containing non-risk allele of SNP rs8062685 (C), 685 RA: promoter vector with enhancer construct containing risk allele (RA) of SNP rs8062685 (G).

## DISCUSSION

Integration of GWAS with multiomic next-generation sequencing techniques offers a powerful approach to decipher GWAS signals in order to implicate both causal variants and their corresponding effector genes. Our prior study focused on the implicated effector gene *PIG-Q* in the modulation of sleep^17^, while this current effort focused on pinpointing the corresponding causal variant.

We leveraged the results from our physical ‘variant-to-gene mapping’ strategy, bringing our attention to three proxy SNPs to the GWAS sentinel signal embedded within the *WDR90* locus. This evidence warranted further investigation to determine if this specific genomic region does indeed exhibit distal enhancer activity for *PIG-Q* in the human cellular context, and if so, to further pinpoint the causal variant(s) among the three proxy SNPs.

*WDR90* knock out iPSCs were generated using a CRISPR-Cas9 approach. The iPSCs were differentiated to neural progenitor cells (NPCs) to investigate changes in the expression of *PIG-Q* in knock-out clones. While these NPCs have a confirmed deletion of the putative enhancer region containing the SNPs, this deletion did not fully eliminate the expression of *WDR90*. The gene-level *PIG-Q* expression level remained unchanged in the knock-out NPCs, but closer appraisal of transcript variants revealed more interesting results. Indeed, it is known that *PIG-Q* has multiple transcript variants and not all are protein coding^24^. Summarizing the RNA sequencing results at the transcript level enabled the identification of transcript specific changes in *PIG-Q*. Two *PIG-Q* transcript variants – *PIG-Q*-208 and *PIG-Q*-218 were found to be differentially expressed. The expression of *PIG-Q*-218 was significantly lower in the knock-out cell lines when compared with the wild type, whereas the expression of *PIG-Q*-208 was significantly increased.

Ensembl^24^ tags *PIG-Q*-218 as TSL5 (Transcript Support Level 5) indicating that there is no evidence or poor evidence of this protein transcript to support the model structure, while *PIG*-Q-208 was tagged as TSL2 (Transcript Support Level 2), with evidence of protein transcript through multiple expressed sequence tags (ESTs)^24^. However, both transcript variants are predicted to be targeted for degradation via NMD, which is a surveillance pathway that exists to decrease errors in gene (*PIG-Q*) expression by eliminating the presence of aberrant mRNAs with premature stop codons^39^. NMD surveillance depends on at least one protein copy being translated from each recognized NMD transcript to determine its fate – to either produce multiple functional outputs of a single pre-mRNA transcript or destroy it to prevent the production of an aberrant protein^40^. While these truncated proteins are generally misfolded and quickly degraded by a proteasome, some could also be stable and functional. Future investigation into these transcript variants is thus warranted for better understanding of *PIG-Q* expression and regulation. Given both differentially expressed *PIG-Q* transcript variants are likely targeted by NMD, we were not motivated to quantify their protein levels by Western blotting.

More than 20 different proteins encoded by Phosphatidyl Inositol Glycan (PIG)/Post GPI Attachment to Proteins (PGAP) gene family are involved in the GPI-anchor biosynthesis. *PIG-Q* is not a crucial component of the N-acetylglucosamine transferase complex but is essential for stabilizing the enzyme complex in the first step of GPI biosynthesis^41,42^. Watanabe *et al.* reported the interaction between *PIG-Q* and other PIG gene products – *PIG-A, PIG-H*, and *PIG-C* ^41^. Hence, we investigated the fate of the *PIG* genes interacting with *PIG-Q* upon deleting the variants that potentially affect *PIG-Q* gene expression. *PIG-A*, *PIG-H* and *PIG-C* were not significantly differentially expressed in the knock-out NPCs, while *PIG-W* was down-regulated in knock-out NPCs; however, the significance of this remains to be elucidated.

Sequencing analysis of the knock-out NPCs showed that they were enriched in events that translated to “intron inclusion” – during splicing, the probability of an intron being included as a *de novo* exon was higher in knock-outs when compared to the unedited NPCs. This result indicates that the CRISPR deletion led to changes in distal gene splicing. At *PIG-Q*, a putative cryptic exon was predicted to be preferentially included in two of the three knockouts.

Putting together these pieces, we hypothesized that the proxy SNPs identified through our variant-to-gene mapping influence the splicing of *PIG-Q* transcripts, rather than its overall expression. All three SNPs are sQTLs for *PIG-Q* in 48 GTEx tissues and eQTLs for *PIG-Q* in 3 brain tissues^43,44^. Recent evidence points to the importance of sQTLS (splicing quantitative trait loci) may play an important role in disease pathogenesis. sQTLs where variants affect alternative splicing may be strong candidates for disease mechanisms as alternative splicing affects gene expression levels^45^. Several studies have demonstrated evidence where alternative splicing is responsible for disease progression^46-48^. Our working hypothesis is that alternative splicing that occurs in the presence of proxy SNPs may alter the ability of *PIG-Q* to bind with other PIG genes and destabilize the GPI-anchor biosynthesis. GPI-anchor biosynthesis is important for many surface expression proteins, and defects in this pathway can result in congenital disorders of glycosylation^49^.

Among the other genes that were significantly different in knock-out NPCs were the genes coding for proteins of the WNT signaling pathway – *RSPO1* and *RSPO3*. Several WNT genes – *WNT4, WNT5B, WNT7A, WNT7B, WNT8B*, and other related genes were down-regulated. The WNT signaling pathway is a key regulator of many growth and developmental processes such as embryogenesis, neurogenesis, etc.^50^. WNT components can impact cell cycle at various checkpoints^51^. The WNT signaling cascade is initiated when a WNT protein ligand binds to a frizzled receptor^52^. Proper functioning of WNTs depend on their regulation through various posttranslational modifications. Recent evidence showed that a subset of WNTs can be modified by (GPI)-anchored metalloproteases, the significance of which is not yet clear^53^.

However, the role of PIG genes, specifically *PIG-Q* in WNT signaling pathway is unclear. Moreover, it remains to be elucidated if the changes in WNT pathway gene expression is due to the 2 kb deletion in the NPCs or the specific removal of the three putative regulatory variants.

Given this confluence of evidence, we concluded that the region harboring these proxy variants is regulatory in nature and influences *PIG-Q* splicing and transcript expression. Thus, we sought to identify which of the three proxy SNPs in LD with the sentinel yield an allelic effect on gene expression. Transcription factor motif analysis for the three proxy SNPs showed that rs3752495 was predicted to affect a putative binding site for transcription factors SP1, EGR1 and EGR2 (Zinc Finger Family) (**Suppl. Methods, Suppl. Fig. 3, Suppl. Table 7**). Therefore, among the three SNPs, rs3752495 was predicted to have the highest probability of being regulatory. Furthermore, the alternative allele (risk allele, C) was predicted to have a higher affinity to the transcription factor binding site than the reference allele (non-risk allele, T) (**Suppl. Table 6**). Our luciferase reporter assays agreed with this *in silico* prediction, with the cells transfected with the putative enhancer region containing the rs3752495 -C (risk allele) showing higher luciferase expression when compared with its non-risk allele as well as other variants.

*PIG-Q* encode a GPI-anchor protein involved in the first step of GPI anchor biosynthesis – *PIG-Q* stabilizes a core complex of *PIG-A, PIG-C* and *PIG-H*^54,55^. The GPI pathway is necessary for posttranslational modification of several proteins involved in membrane trafficking to facilitate several cell processes including neurogenesis^56,57^. *PIG-Q* is expressed in the brain, and its homozygous deletion was previously found to be lethal in a mouse model^58^. In humans, rare exonic mutations in *PIG-Q* are known to cause syndromic forms of epilepsy and epileptic encephalopathy^18,59,60^ but its role in sleep and sleep disorders is not yet clear. Many sleep-associated loci are pleiotropic and may influence sleep either directly or indirectly via their associations to other disorders including epilepsy, metabolic, or neuropsychiatric dysfunction (e.g., *MEIS1*). Several Mendelian randomization studies have shown a significant association between sleep and neuropsychiatric disorders^61^. Given this complex relationship, we cannot ignore the potential involvement of a *PIG-Q*/epilepsy connection in modulating sleep. How this gene’s mechanism of action potentially bridges sleep and seizures, therefore, requires further study.

## Limitations

While one of the three candidate SNPs showed a regulatory effect by luciferase assay, this experiment design utilized a generic SV40 promoter so we cannot conclude if genotype at rs3752495 directly regulates *PIG-Q* expression *in vivo*. We also identified no major differences in the expression of *PIG-Q* at the gene level; however, we did observe differences in individual transcript abundances and a significant difference in one other PIG gene. The knock-out NPCs and the unedited NPCs used for RNA-seq analysis were generated from iPSCs at different passage numbers. Although all cell lines used for differentiation had a normal karyotype, differences in culture conditions, although minor, could potentially lead to some variations in our observations and conclusions. The rationale behind generating knock-outs instead of making base alterations was based on the expectation that a gross genomic change would yield more obvious differential gene expression of a higher magnitude thus motivating to do more discrete work subsequently, rather than a single base change at a subtle susceptibility variant; however, this was not entirely the case in our study. We did not carry out experiments to validate the binding of predicted transcription factors such as ChIP-seq or electrophoretic mobility shift assay (EMSA). Since the causal variant is in a transcription factor binding motif, ChIP or EMSA could verify predicted differences in transcription factor binding in the presence of risk allele and further characterize the pathway by which this variant likely contributes to insomnia. Finally, we cannot rule out that the pathway expression effects are at least in part influenced by the direct perturbation of *WDR90* expression, although we have no evidence of this gene’s relationship with sleep or the variants of interest.

## Conclusion

In this study, we aimed to further characterize an insomnia GWAS locus in order to understand the relationship between the candidate causal variant(s) within the *WDR90* locus and their role in modulation of *PIG-Q*. Our variant-to-function approach and subsequent *in vitro* validation implicates rs3752495 as a causal insomnia risk variant embedded within a *WDR90* intron influencing *PIG-Q* splicing. Future use of mammalian animal models could aid in further understanding of this relationship. Development of a humanized mouse model harboring the *WDR90* embedded variants identified through GWAS could reveal how these variants affect the expression and/or function of *PIG-Q* in an *in vivo* setting. Both *PIG-Q* and rs3752495 represent promising insomnia intervention points, but further functional studies are required to investigate the role of *PIG-Q* and GPI-anchor biosynthesis as a whole in sleep/wake regulation.

## ACKNOWLEDGMENTS

This work was in part supported by National Institutes of Health award R01 HL143790 and the Daniel B. Burke Endowed Chair for Diabetes Research. S.S. designed the study, collected and analyzed data, wrote and edited manuscript; S.H.L. analyzed data, wrote/edited manuscript; M.C.P. analyzed data, wrote/edited manuscript; A.J.Z. analyzed data, reviewed and edited manuscript; A.C. analyzed data, reviewed and edited manuscript. C.L. analyzed data, reviewed and edited manuscript; E.B. reviewed and edited manuscript; J.A.P collected and analyzed data, reviewed and edited manuscript; A.D.W. reviewed and edited manuscript; F.D.B. reviewed and edited manuscript; A.I.P. reviewed and edited manuscript; P.R.G. reviewed and edited manuscript; A.C.K. reviewed and edited manuscript; S.F.A.G. supervised the study, reviewed and edited manuscript. Sonti and Grant are the guarantors of this work and, as such, had full access to all the data in the study and takes responsibility for the of the data and the accuracy of the data analysis.

## Supplementary Figure legends

**Supplementary Figure 1:**
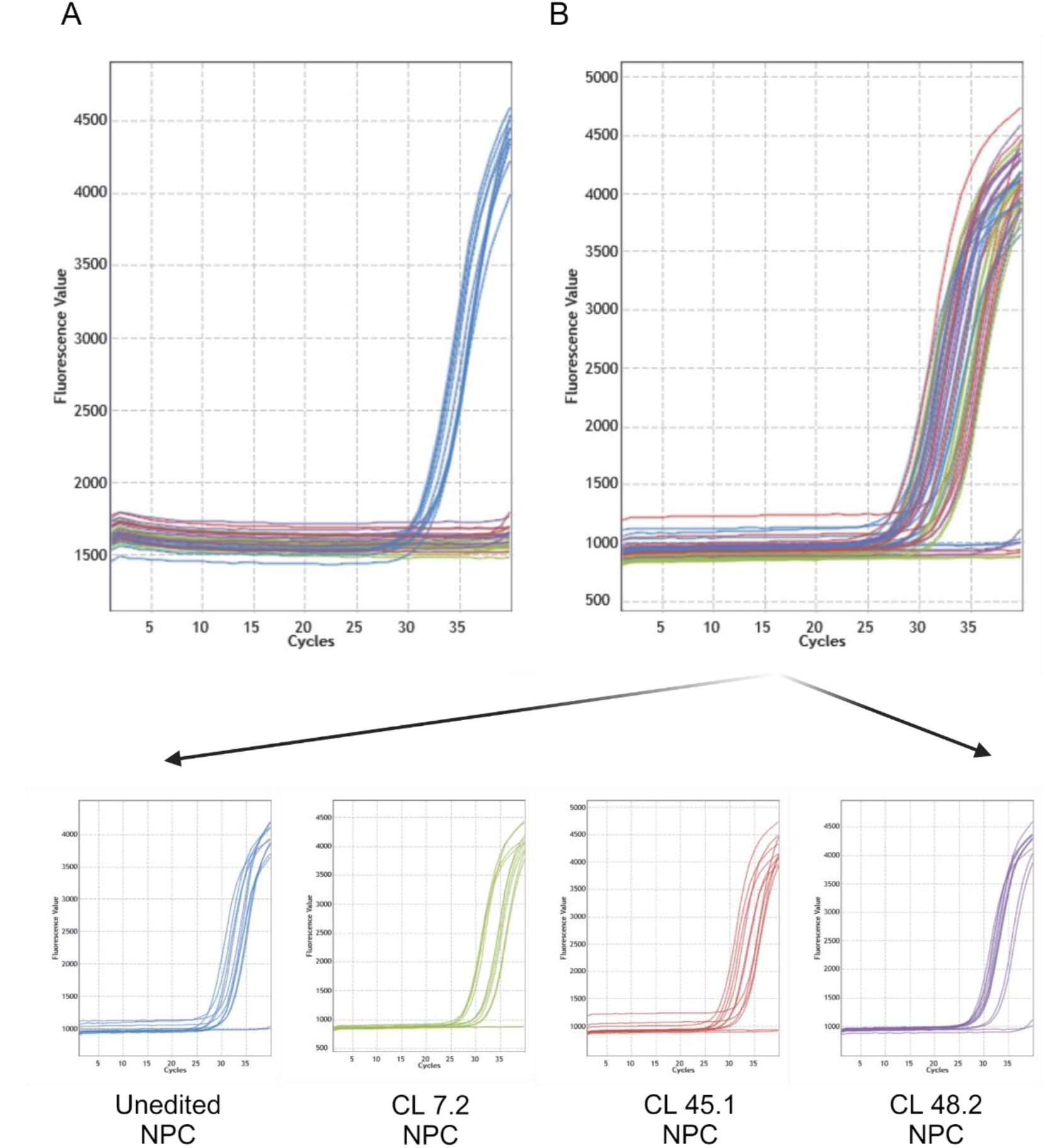
The deletion within *WDR90* was confirmed with PCR. **a.** Amplification plot of the CRISPR deleted region in *WDR90*: Amplification was observed only in unedited NPCs but not in edited knock-out NPCs. b. Amplification plot of the untargeted region in *WDR90*; Amplification was observed both in unedited NPCs as well as edited knock-out NPCs. (Blue, unedited NPCs; red, KO CL 7.2; green, KO CL 45.1; purple, KO CL 48.2)

**Supplementary Figure 2:**
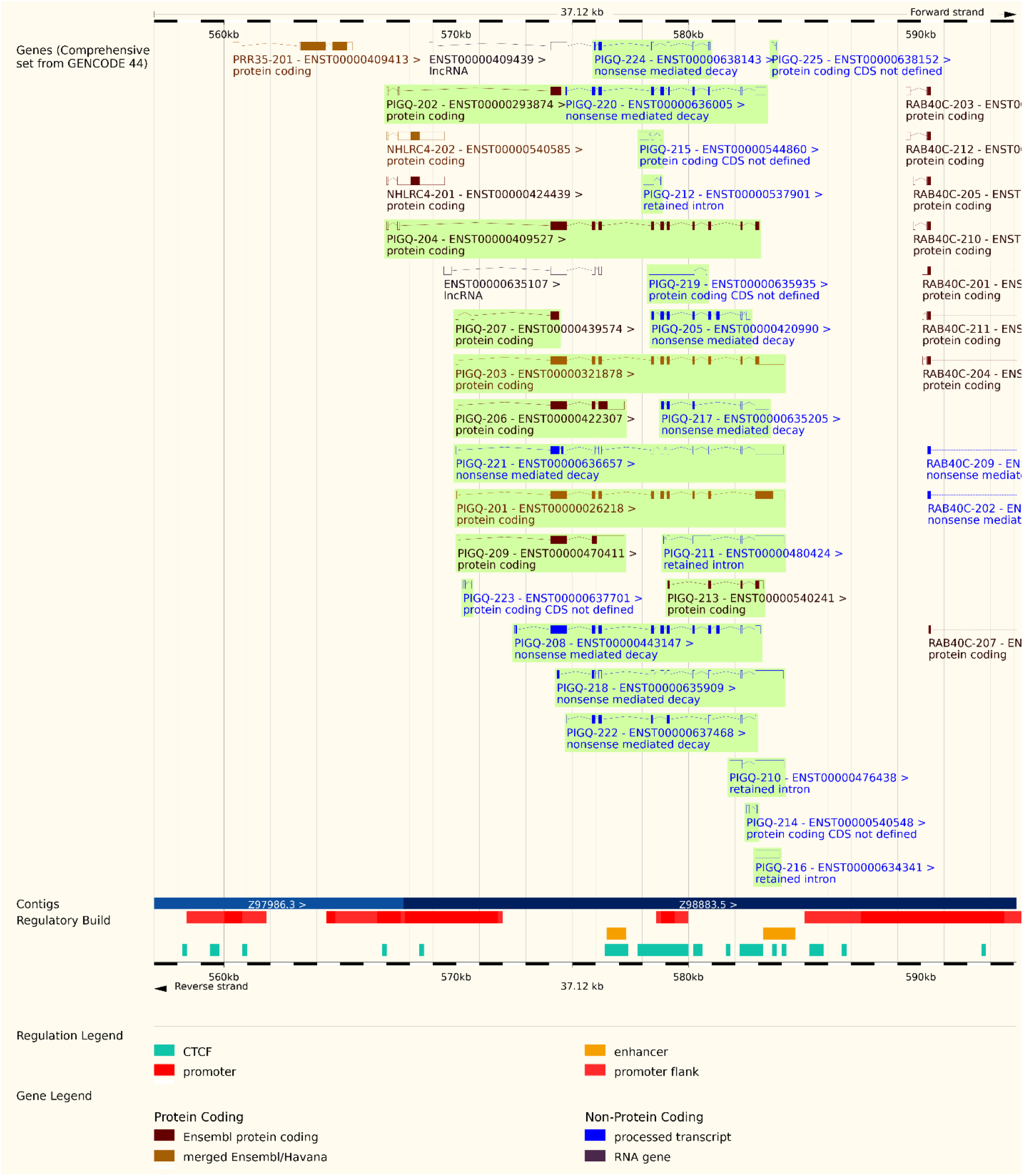
*PIG-Q* transcript variants. Representation of *PIG*-Q transcript variants as reported on Ensembl.

**Supplementary Figure 3:**
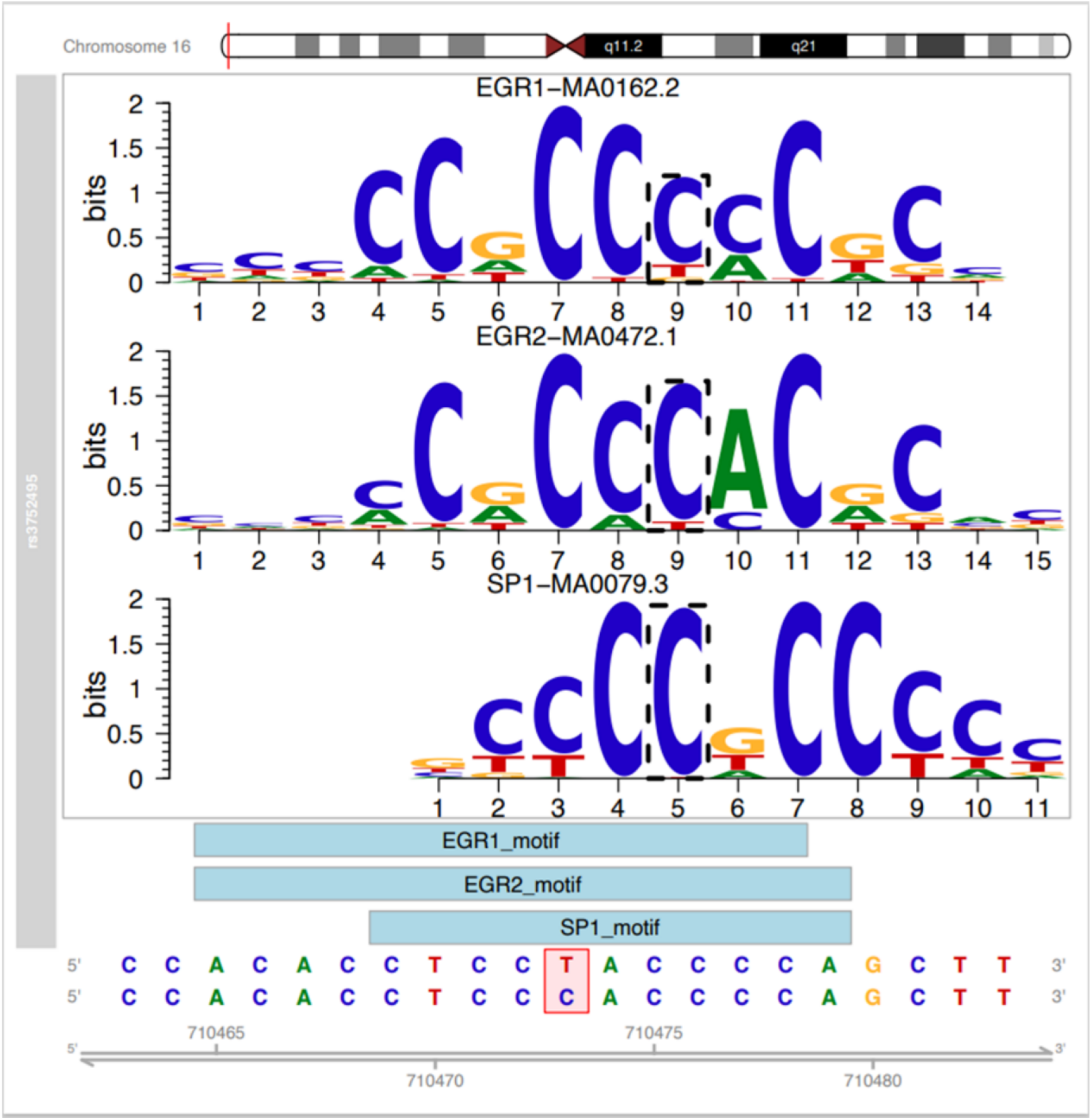
Transcription factor Motif analysis.

